# Perineuronal nets in the bed nucleus of stria terminalis mediate stress-induced anxiety behaviour in adult male mice

**DOI:** 10.64898/2026.10.15.759345

**Authors:** Kathryn Murrall, Mariusz Mucha, Kratika Mujmer, Marc Ten Blanco, Ben Sherlock, Robert Pawlak, Valentina Mosienko

## Abstract

Anxiety disorders affect an estimated 301.4 million people globally. However, they remain among the most challenging conditions to treat, with only around half of patients reporting a 50% or greater reduction in their symptoms. The discovery of novel and more effective treatments therefore requires a better understanding of the underlying mechanisms of anxiety at both the circuit and cellular levels.

Stress induces aberrant changes in neuronal plasticity across brain regions within the anxiety circuitry including the bed nucleus of the stria terminalis (BNST). One mechanism through which synaptic plasticity is regulated is the organisation of perineuronal nets (PNNs), specialised extracellular matrix structures that are highly sensitive to stress. In rodent models, stress is associated with a reduction in PNN number and/or decreased *Wisteria floribunda* Agglutinin (WFA) labelling in limbic areas. However, the development of PNNs within the BNST and their role in the stress responsiveness remain largely unknown.

In the current study we investigated the spatiotemporal development of PNNs within the BNST of male mice, where they first appear at P30 and continue to mature into adulthood, reaching their highest expression at P210 and specifically within the anteromedial BNST. We further showed that stress-induced anxiety is associated with a reduction in PNN number and significant changes to PNN fine structure including increased leakiness in the anteromedial BNST. Most importantly, enzymatic removal of PNNs by chondroitinase ABC in the mouse anteromedial BNST exacerbated behavioural responses to acute stress as demonstrated by increased anxiety-like behaviour. Collectively, these results highlight an indispensable role for PNNs within the anteromedial BNST in mediating a healthy emotional response to psychosocial stress.

## INTRODUCTION

Anxiety disorders are the most prevalent psychiatric conditions worldwide, affecting an estimated 301.4 million people. Anxiety disorders are pervasive, often chronic and the leading cause of disability globally [1]. Nevertheless, anxiety disorders remain some of the most difficult conditions to treat. Response to treatment for anxiety disorders is highly variable, with only about half of patients reporting at least a 50% reduction in their symptoms [2, 3]. To identify novel and effective treatments for anxiety, it is essential to better understand its underlying mechanisms at both the circuit and cellular levels.

Stress is thought to promote anxiety through maladaptive plasticity in the prefrontal cortex (PFC), hippocampus and amygdala [4–6]. The bed nucleus of the stria terminalis (BNST) is a further key node of this circuit, integrating limbic inputs and projecting to hypothalamic and brainstem regions that control stress responses [7, 8]. BNST lesions attenuate anxiety-like behaviour and disrupt cortisol release in animal models [9–12]. The BNST is also engaged during sustained threat in both rodents and human [7, 8, 13, 14]. However, the direction of the response depends on the microcircuitry originating in a specific BNST subnuclei. For example, the oval nucleus promotes anxiety, whereas the anterodorsal BNST reduces it [8, 15]. The anteromedial BNST (STMA) is less well studied, but its inputs from the medial and basomedial amygdala and the ventral subiculum suggest that it contributes to anxiety responses to threatening odours and contexts [16–18].

Persistent alterations in neural network plasticity are among the likely mechanisms underpinning pathophysiological anxiety [19–21]. One important regulator of synaptic plasticity is the extracellular matrix (ECM), which in some parts of the brain condenses into uniquely organised perineuronal nets (PNNs) around neuronal bodies and/or dendrites [22]. PNNs form at the end of critical periods and limit plasticity in the adult brain, while their removal in adulthood with chondroitinase ABC (ChABC) restores juvenile-like plasticity [23]. In the amygdala, PNNs make fear memories resistant to erasure [24], pointing to a role in stabilising emotional memory. PNNs nevertheless remain dynamic throughout life [25], and their remodelling can reopen windows of plasticity.

PNNs are sensitive to stress. However, the direction of stress-triggered PNN remodelling depends on the brain region, the type and duration of the stressor, the developmental stage at which it is experienced, and the time between stress and PNN examination. Reductions in PNN number or labelling have been reported after chronic restraint stress in the medial PFC [26, 27], after maternal separation in the infralimbic cortex [28] and after early-life adversity in the ventral hippocampus [29]. In contrast, increase in PNN density or labelling have been described in the hippocampal CA1 after chronic social defeat [30], in the mPFC after chronic variable stress [31, 32] and in the right basolateral amygdala after early-life stress [33].

PNNs have been detected previously in the BNST [34–36]. However, their development across subnuclei, response to stress and role in anxiety are unknown. Because the BNST continues to mature into adulthood [37], PNNs here may develop later than in other limbic regions. We hypothesised that PNNs in the anteromedial BNST stabilise circuits that regulate emotional responses, so that intact nets limit stress-induced anxiety while their degradation weakens this protection. If so, stress should reduce the number and structural integrity of BNST PNNs, and removing them should worsen anxiety-like behaviour after stress.

To test this, we mapped the spatiotemporal development of PNNs across BNST subnuclei from early postnatal life, quantified the effects of repeated restraint stress on PNN number and fine structure, and examined the effect of ChABC-mediated PNN removal on stress-induced anxiety-like behaviour.

## METHODS

### Animals

All procedures were carried out in accordance with the UK Animal (Scientific Procedures) Act 1986. Experiments were approved by the UK Home Office (P5EAA607F, PP5495972) and the University of Exeter and Bristol Local Ethics Committees. Male C57BL/6J mice (Charles River, UK) were group housed, in temperature-controlled rooms, on a 12-hour light/dark cycle, with lights on at 6:30am, with ad libitum access to food (Teklad Global 14, 14% protein diet, Envigo) and water.

### Stereotaxic surgery

Mice were first anaesthetised in a chamber with isofluorane (2-3%) and oxygen (2 L/min), and then transferred to a gas mask, where isofluorane (1-1.5%) and oxygen (1L/min) continued, for surgical preparation. Mice were shaved and cleaned with Hibiscrub from the neck to behind the eyes, and given pre-surgery analgesia intraperitoneally (0.15mg/kg buprenorphine or 0.5mg/ml Rimadyl) before transfer to the stereotaxic frame. Depth of anaesthesia was monitored by testing the pedal reflex whilst the mouse was using the gas mask and following transfer to the stereotaxic frame.

Before making an incision on the skull, stray hair were removed and the skin was treated with EMLA (Eutectic Mixture of Local Anesthetics, 2.5% prilocaine and 2.5% lidocaine) for 5 minutes, then was cleaned and sterilised using Betadine. Sterility of the surgical area was maintained throughout the procedure. After a skull incision in the skin handheld dental drill and sterile 0.5mm diameter carbide drill burrs (World Precision Instruments, 501860) were used to create entry-points for injections into the brain with any debris or bone fragments being cleared before beginning injections.

Anteromedial BNST injections were made bilaterally at 0.38 mm anteroposterior, +/- 0.5 mm mediolateral and -3.75 mm dorsoventral to bregma. ChABC enzyme (Sigma Aldrich, UK; 50U, 300nl/site), or sterile PBS (0.5 M; Sigma Aldrich, UK) was injected at a rate of 100-250nl per minute using a 35G bevelled needle (World Precision Instruments, USA) inserted into a 10 μL Hamilton syringe connected to a microsyringe pump.

Immediately following surgery, each animal was transferred to a cage with a heating mat to ensure temperature regulation and monitored for 45-60 minutes post-surgery before being returned to their home cages. Surgery animals were given DietGel® Recovery (ClearH20®, 72-06-5022), HydroGel® (ClearH20®, 70-01-5022) and a mash made from standard animal food pellets ad libitum and their weight gain was monitored for 5 days after surgery.

### Labelling and imaging of perineuronal nets

10, 30, 60, 210 or 365 days old mice (spatiotemporal development of PNNs) or adult, 10-12 week old stress naïve (control) animals or mice that underwent restraint stress (stress) were anaesthetised with sodium pentobarbital (50 mg kg-1) and transcardially perfused with ice-cold phosphate buffered saline (PBS; Sigma) followed by room temperature paraformaldehyde (PFA, 4% in PBS; Sigma). Following dissection and overnight fixation with 4% PFA at 4°C, brains were washed in PBS, transferred to a cryoprotective solution (30% sucrose in PBS) for dehydration, and stored at 4°C until sectioning Sequential 30 μm coronal anterior BNST sections, bregma 0.62 mm to 0.02 mm [70], were cut on a cryostat (Bright Instruments, UK), with every third section being taken then for PNN labelling.

PNNs within the BNST were identified using *Wisteria floribunda* agglutinin (WFA), a lectin that selectively binds N-acetylgalactosamine residues on glycoproteins, and is widely used as a marker for perineuronal nets in the brain [29, 41, 43, 71–74]. Following incubation in blocking solution (10% foetal bovine serum; Sigma, UK in 0.1% PBST: Triton X-100 in PBS; Sigma, UK), sections were incubated in FITC conjugated WFA at room temperature (1:500; VectorLabs, UK) in blocking solution, and following wash mounted on Poly-D-Lysine coated slides (VWR, UK) with mounting medium (Fluorsave Merck, UK) and imaged on a Zeiss LSM 5 Exciter confocal microscope (Zeiss, Germany). The Franklin and Paxinos mouse brain atlas was used as a guide to create masks of bregma points throughout the BNST, from approximately bregma 0.62 mm to 0.02 mm [70]. Masks were made transparent and were overlaid onto microscopy images of the BNST in accordance with bregma point, using the anterior commissure as an anatomical reference guide, to separate the distinct subnuclei.

### Evaluation of perineuronal net number and intensity

For manual counting of PNNs, images with mask overlays were opened in ImageJ (v1.53e; NIH, USA) and the count tool was used to identify PNN positive cells. PNN identification was based on the criteria that they must be independently visible and distinctly reticular. Meshes of PNN units were not counted, as individual units could not be separated. Intensity of PNNs was measured using a MATLAB (Mathworks, USA) script (Supplementary information) which parcellated each distinct BNST subnucleus. To account for background fluorescence, the average pixel intensity of a separate region, adjacent to the BNST, where no PNNs were expressed, was subtracted from the intensity measured within BNST regions on the same slice. As absolute scale was not factored into the automated analysis, values are relative and presented in arbitrary units.

### Quantification of perineuronal net fine morphology

PNN fine morphology was analysed from three-layer z-stack magnified images with 0.1 μm between stacks with the second layer of the stack being of optimum focus on the PNN surface, using APNU v1.1 Image J plugin as previously described [75]. Briefly, after manual selection of the central point within each hole of an individual PNN unit, the programme calculated an outline of the PNN from the three confocal layer images. The output text file from the script provided values for a number of distinct parameters including: hole area (Area3Dhole), hole perimeter (Per3DECM) and intensity of matrix surrounding the hole (IntMean3DECM). In addition, PNN flatness was measured by the standard deviation of ECM intensity across the three Z planes (zSDECM), and PNN leakiness was measured by the minimum intensity of ECM in 3D space (IntMin3DECM).

### Repeated acute restraint stress model

10-12 weeks-old mice were restrained for 2 hours (10am-12pm) for 3 consecutive days in ventilated 50 mL polypropylene conical tubes (Starlab, UK) placed in their home cage. Following restraint mice were immediately released back into the home cage, and tubes were cleaned with soapy water and then ethanol, and reused in subsequent experiments. Control animals were left undisturbed in their home cages until the next phase of the experiment.

### Elevated plus maze

The apparatus was fabricated from melamine coated medium-density fibreboard (Technical Services, University of Exeter Mechanical Workshop) and consisted of two open arms (30 cm in length) and two closed arms (30 cm in length, with 12 cm high walls), with a 10 cm × 10 cm central square, elevated 60 cm off the floor on metal legs. The open arms of the maze were brightly lit for the duration of the experiment (254 lux). Mice were individually placed in the centre square of the maze, facing an open arm, and were allowed to freely explore for five minutes. Behaviour was video recorded (Logitech, Switzerland) and subsequently analysed using Viewer2 (Biobserve GmbH, Germany) recording number of entries and time spent in each zone.

### Statistical analysis

Graphpad Prism (Graphpad Software Inc, USA) was used to perform all statistical analyses. Unpaired t-test or one-way ANOVA with appropriate post-hoc test (indicated in figure legends) was used to determine significant differences between group means. Data are presented as mean ± SD unless otherwise stated.

## RESULTS

### Perineuronal nets in the bed nucleus of the stria terminalis are detected at postnatal day 30 with the highest density in the anterior part of the medial division

To investigate the development of PNNs in the BNST over time, we labelled BNST sections from mice aged postnatal days (P) 10, 20, 30, 60, 210 and 365 with WFA. These time points were chosen to represent successive life stages of the mouse, which also correspond to human life stages. The two early time points, P10 and P20, represent early life, P20 in particular was chosen as PNNs have been identified in other brain regions by this time [38–40]; P30 represents early adolescence; P60, late adolescence; P210, mature adulthood and finally P365, middle age [25].

No PNNs were detected in the BNST at P10 (Figure 1). By P20 WFA labelling was detected revealing that the matrix was beginning to condense, as shown by a greater intensity of the labelling, though no evident PNNs were yet identified. The characteristic shape of PNNs became apparent in the BNST at P30, with more intense WFA labelling around neuronal cell bodies and proximal dendrites (Figure 1). At P60, PNNs in the BNST were visually similar to those documented in other brain regions, including the hippocampus and amygdala (Figure 1; [38, 39]). Notably, PNN structure in the BNST is less reticular, lacking the distinct lattice-like structure observed in PNNs in the cortex and hippocampus, and more granular, like PNNs in the basolateral amygdala. PNNs in the BNST do not show the same organisation pattern as in the cortex, hippocampus, or amygdala. Instead, they are more densely clustered together in a small region, in contrast to the few, evenly spaced PNNs in the amygdala, the random, widespread expression of PNNs in the cortex and the highly organised PNNs in the hippocampus.

**Figure 1.**
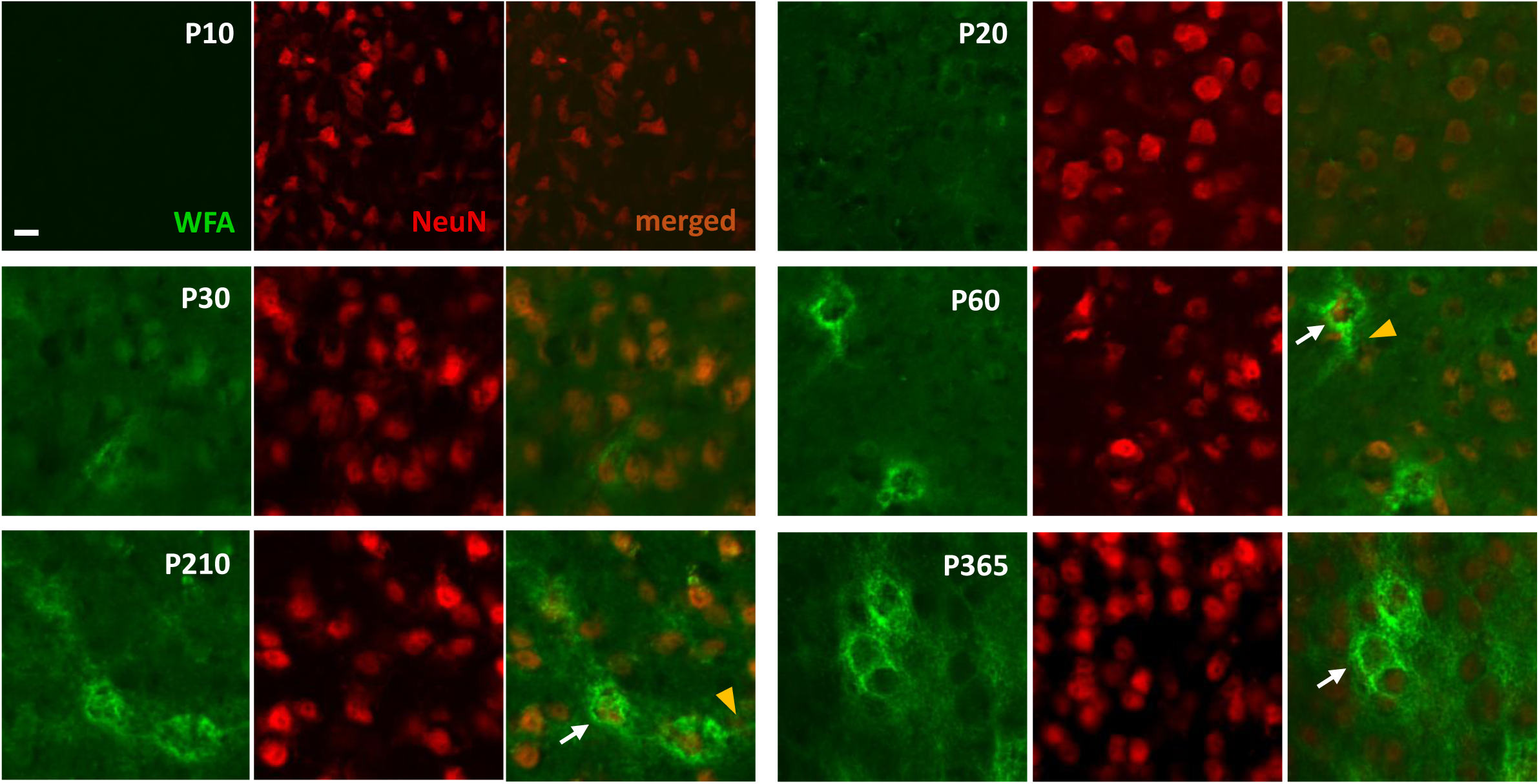
Temporal development of perineuronal nets in the bed nucleus of the stria terminalis. WFA labelling (green) in the bed nucleus of the stria terminalis (BNST) is undetectable at postnatal day (P) 10. Weak WFA labelling can be detected at P20, and by P30, the characteristic ‘net’ shape can be identified around neuronal cell bodies labelled with anti-neuN antibodies (red). WFA labelling is stronger from P60 and is detected around both neuronal cell bodies (white arrows) and their proximal dendrites (orange arrows) in the BNST, with most nets being clustered together. Scale bar = 20µm. WFA labelling of BNST slices was repeated in 5-6 naïve mice for each time point, where all qualitatively showed the same result.

PNNs were also visualised at two much later points in life: P210 and P365. PNNs have previously been shown to be developmental features important for the closure of the critical period of synaptic plasticity [23, 41, 42]. However, PNNs are also proposed to undergo remodelling throughout adulthood, based on activity in local microcircuits [43]. WFA labelling revealed that at P210 and P365 PNNs in the BNST are more densely expressed than in early life, and in some instances form clusters around adjacent neurons (Figure 1).

To quantify the development of PNNs in the BNST, we employed both manual and automatic methods. The P10 and P20 time points were excluded from quantitative analysis of PNN spatiotemporal expression since no PNNs were detected in mouse brains at these time points (Figure 1).

We first counted the number of PNNs manually using ImageJ. PNNs were expressed in the STMA at P30, and across the BNST from P60 becoming more numerous and intense throughout development (Figure 2). PNN number also increased significantly with age in the STMA (Figure 3A; F(3,18) = 16.48, P = <0.0001). The number of PNNs doubled between P30 and P60, and between P60 and P210 (Figure 3A; P30: 6.10 ± 3.47 vs P60: 16.59 ± 8.25, p = 0.0911; P60 vs. P210: 31.42 ± 8.61, p = 0.0079). There was no difference in PNN number between P210 and P365 (Figure 3A; P365: 28.00 ± 4.28). PNN number increased significantly over time in the STMV (Figure 3A; F(3, 18) = 3.787, P = 0.0288). Specifically, a greater number of PNNs was expressed at P365 compared with P30 (Figure 3A; P30: 4.50 ± 4.5 vs. P365: 12.05 ± 3.71, p = 0.0198). No difference in PNN number was observed in the STMV at P60 or P210 compared to P30 (P60: 8.55 ± 4.19; P210: 10.02 ± 2.85). PNN number did not change over time in the STLV (Figure 3A; P30: 5.70 ± 3.74; P60: 7.95 ± 5.00; P210: 8.13 ± 4.01; P365: 6.67 ± 3.71). In both lateral subnuclei of the BNST, (STLP and STLD), the number of PNNs increased throughout development. In the STLP, PNN expression increased with age (Figure 3A; F(3, 18) = 11.96, P = 0.0002). Specifically, more PNNs were expressed at P210 than at the P30 or P60 time points (Figure 3A; P210: 9.67 ± 2.16 vs. P30: 3.9 ± 1.34, p = 0.0002 and vs. P60: 6.5 ± 2.29, p = 0.0360). Twice as many PNNs were expressed in the STLP at P365 compared with P30 (Figure 3A; P30 vs. P365: 8.88 ± 0.80, p = 0.0009).

**Figure 2.**
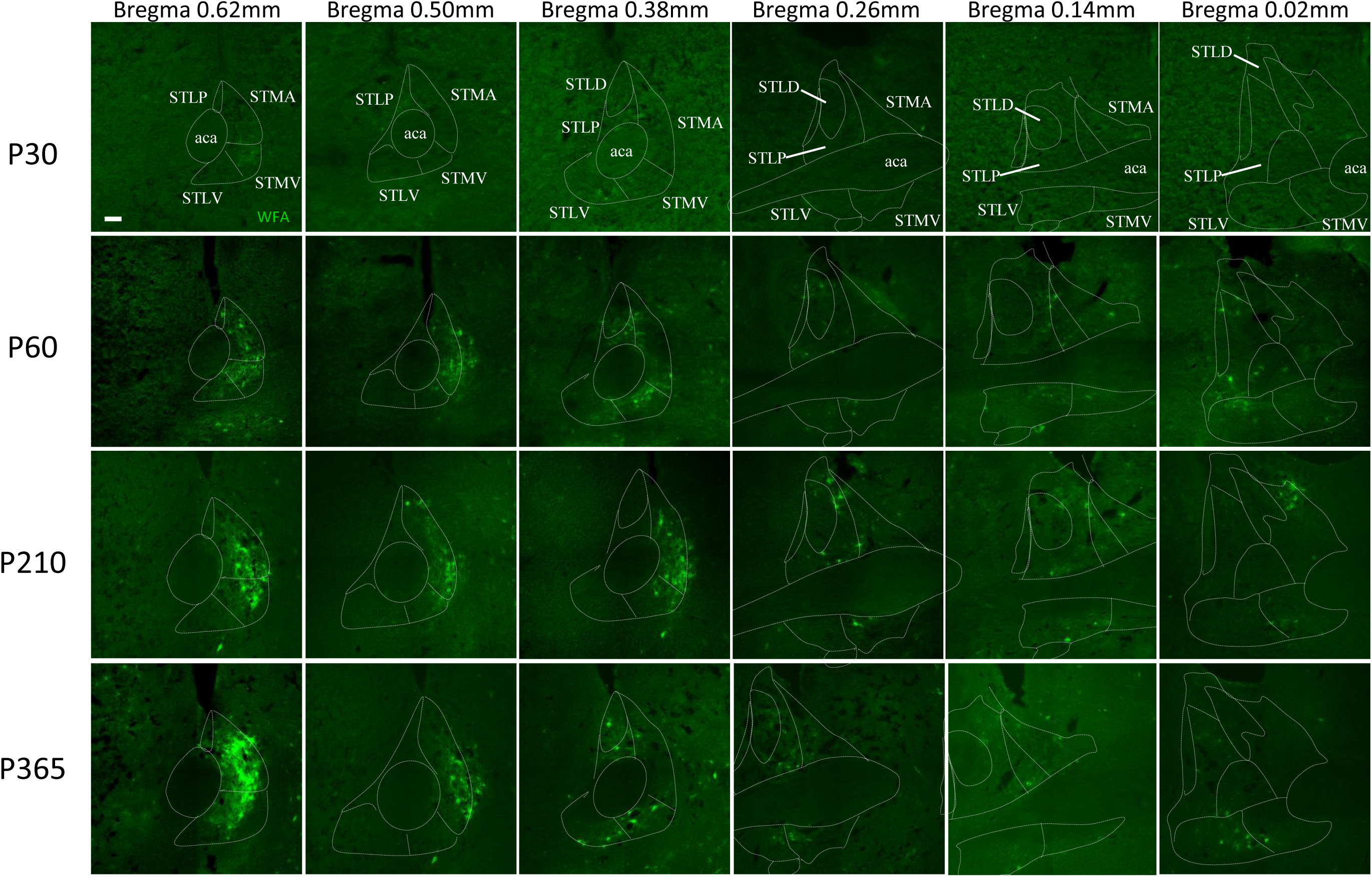
Spatiotemporal distribution of perineuronal nets in the bed nucleus of the stria terminalis. WFA-labelled perineuronal nets (PNNs, green) show a similar spatial distribution within the bed nucleus of the stria terminalis (BNST) throughout development, from P30 to adulthood (P60), and beyond (P210 and P365). PNNs expression is stronger in the medial subnuclei in rostral parts, and the dorsomedial and ventromedial nuclei in caudal parts of the BNST. Masks were created using mouse brain atlas (Franklin & Paxinos, 2007) and indicate the largest BNST subnuclei of interest. Abbreviations: STMV – BNST, medial division, ventral part; STLV – BNST, lateral division, ventral part; STMA – BNST, medial division, anterior part; STLP – BNST, lateral division, posterior part; STLD – BNST, lateral division, dorsal part; aca – anterior commissure. Scale bar = 100µm. WFA labelling of BNST slices was repeated in 5-6 naïve mice for each time point, where all qualitatively showed the same result.

**Figure 3.**
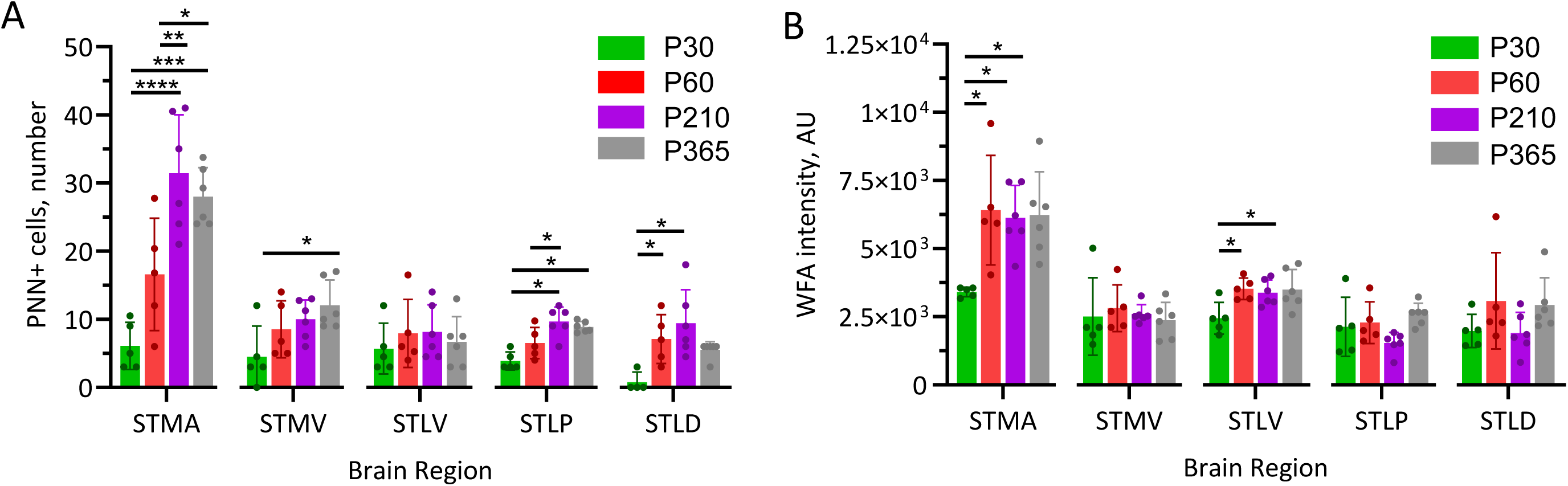
Quantification of perineuronal nets within anterior subnuclei of the bed nucleus of the stria terminalis throughout development. Perineuronal net (PNN) number (A) and intensity of WFA labelling (B) were quantified through development in the medial division, anterior (STMA) and ventral (STMV) parts, in the lateral division, ventral (STLV) and posterior (STLP) parts, and in the lateral division 2 (STLD) of the bed nucleus of the stria terminalis (BNST). Data are presented as bar plots with individual values and mean±SD; N (P30/P60) = 5 mice, N (P210/P365) = 6 mice, 6 brain sections per animal, PNN number/intensity averaged across left and right hemispheres. *p < 0.05, **p < 0.01, ***p < 0.001, ****p < 0.0001 vs an indicated group, one-way ANOVA with Tukey’s post hoc testing.

In the STLD, the number of PNNs significantly increased with age (Figure 3A; F(3, 18) = 7.102, P = 0.0024). There were seven times more PNNs in the STLD at P60 compared to P30 (Figure 3A; P30: 0.60 ± 1.34 vs. P60: 7.10 ± 3.58, p = 0.0244; ). There was no difference in the number of PNNs detected in the STLD at P210 or P365 compared with any other time point (Figure 3A; P210: 9.41 ± 4.95; P365: 5.50 ± 1.23). Overall, PNNs increased in number with age across various BNST subnuclei (Figure 3A). However, there were also a differences in PNN expression throughout development across BNST subnuclei, with the anteromedial part of the BNST (STMA) expressing greater numbers of PNNs than the STLP in adulthood (P60) (Supplementary Table 2; F(4, 20) = 3.312, P = 0.0309; STLP: 6.51 ± 2.29 vs. STMA: 16.59 ± 8.25, p = 0.0366). At P210, the STMA expressed at least three times more PNNs than any other subnucleus (Supplementary Table 3; F(4, 25) = 23.15, P = <0.0001; STMA: 31.42 ± 8.61 vs. STMV: 10.02 ± 2.85, p = <0.0001; vs. STLV: 8.13 ± 4.01, p = <0.0001; vs. STLP: 9.67 ± 2.16, p = <0.0001; vs. STLD: 9.42 ± 4.95, p = <0.0001; vs. ). See Supplementary Tables 1-4 for detailed comparisons of PNN number across BNST subnuclei from P30 toP365.

We also measured PNN intensity computationally using a custom MATLAB script (Supplementary methods). This method provided the advantage of unbiased and automated quantification of PNNs within various BNST subnuclei with the possibility of analysing numerous images at once. PNN intensity increased over time, though not in all BNST subnuclei when quantified using the automated analysis pipeline (Figure 3B). In the STMA, WFA labelling of PNNs exhibited greater intensity throughout development (Figure 3B; F(3, 18) = 5.245, P = 0.0089). WFA labelling was twice as intense at P60, P210 and P365 compared to P30 (Figure 3B; P30: 3411 ± 178 vs. P60: 6413 ± 2011, p = 0.0166; vs. P210: 6127 ± 1191, p = 0.0244; vs. P365: 6234 ± 1579, p = 0.0188). In the STMV and both lateral subnuclei of the BNST (STLP and STLD) no change in WFA labelling intensity was detected (Figure 3B; STMV P30: 2508 ± 1424; P60: 2814 ± 858; P210: 2596 ± 345; P365: 2365 ± 662; STLP P30: 2126 ± 1083; P60: 2283 ± 768; P210: 1534 ± 389; P365: 2589 ± 406; STLD P30: 1977 ± 612; P60: 3082 ± 1763; P210: 1905 ± 753; P365: 2931 ± 1005). Labelling of PNNs in the STLV increased in intensity with age (F(3, 18) = 4.263, P = 0.0193). More intense WFA labelling within the STLV was observed at P60 and P365, compared to P30 (P30: 2443 ± 581 vs P60: 3523 ± 398, p = 0.0342; vs. P365: 3500 ± 731, p = 0.0298). No difference in WFA intensity was observed at P210 compared to other ages (P210: 3381 ± 472). Automated analysis confirmed the greatest PNN expression, by intensity, in the STMA in adulthood (P60) (Supplementary Table 6; STMA: 6413 ± 2011 vs. STLV: 3523 ± 398, p = 0.0179; vs. STMV: 2814 ± 858, p = 0.0027; vs. STLP: 2283 ± 768, p= 0.0006; vs. STLD: 3082 ± 1763, p = 0.0055). See Supplementary Tables 5-8 for comparisons of WFA labelling intensity across BNST subnuclei at P30, P210 and P365, respectively.

In summary, we quantified the spatiotemporal development of PNNs within the BNST. PNNs at first were detected in the BNST at P30, and continue to develop into adulthood, with the greatest expression at P210 in several BNST subnuclei. The subnucleus with the greatest number and intensity of PNNs across developmental stages was the anteromedial BNST, we therefore focused our further studies on this region.

### Stress-induced anxiety is accompanied by a reduced number of perineuronal nets in the bed nucleus of the stria terminalis

We next investigated whether a repeated acute restraint stress paradigm would increase anxiety-like behaviour in mice. The restraint protocol consisted of three consecutive days of 2-hour restraint, and behaviour in the elevated plus maze (EPM) was measured 7 days after the final session (Figure 4A). Repeated acute restraint stress increased anxiety-like behaviour in mice as measured by a two-fold decrease in proportion of open-arm entries (Figure 4B, control: 39.6 % ± 7.3 %, stress: 21.6 % ± 11.3 %, p=0.0016). Following stress, mice exhibited a threefold decrease in both the frequency of open arm entries (stress: 2.3 ± 1.5 entries vs. control: 7.0 ± 2.8 entries, p=0.001) and the time spent in the open arms (stress: 17.4 s ± 12.6 s vs. control: 59.7 s ± 21.5 s, p=0.0003) compared to control mice, both parameters indicative of increased anxiety-like behaviour following stress. However, since stress significantly affected locomotor activity (Figure 4B; control: 9.43 m ± 1.66 m, 7 days post-stress: 6.89 m ± 2.50 m, p=0.0390), the proportion of entries to open arms was used here as a measure of avoidanceand anxiety-like behaviour on the maze.

**Figure 4.**
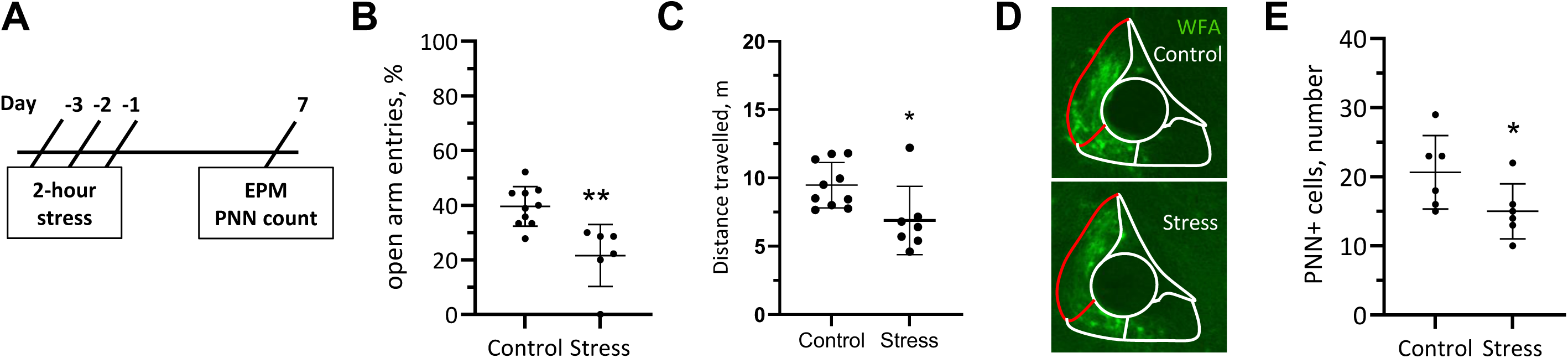
Stress-induced anxiety is accompanied by the reduced number of perineuronal nets in the bed nucleus of the stria terminalis. (A) Schematic timeline of the experiment: behavior on the elevated plus maze (EPM) and perineuronal net (PNN) number were analyzed 7 days ollowing the exposure to 2-hour restraint stress performed during 3 consecutive days. (B) Repeated acute restraint stress increased anxiety-like behavior in mice, as measured by a decreased proportion of open-arm entries on the EPM, and decreased locomotor activity, measured as distance traveled on the maze. (D) Representative images of WFA-staining within the anterior part of the medial division of the bed nucleus of the stria terminalis (STMA, outlined in red line) in stress naïve (control) animals and mice exposed to restraint stress (stress). (E) Number of PNNs in the STMA is reduced following stress. Data are presented as individual values and mean±SD; B: N (control) = 10; N (stress) = 6; D: N(control/stress, animals) = 6, 3 sections per animal, PNN number averaged across left and right hemispheres; *p<0.05, **p<0.01, unpaired t-test

Next, we quantified the effects of acute repeated stress on PNN number. PNN count in the STMA was reduced by 25% following repeated restraint stress (Figure 4C-D, stress: 15 ± 1.6 count vs. control: 20.7 ± 2.2 count, p = 0.039).

Overall, these experimental findings uncover a relationship between PNNs in the STMA and the stress response. Stress-induced increase in anxiety-like behaviour in the EPM is accompanied by a reduction in PNN number.

### Perineuronal net hole area and perimeter increase, and nets show greater leakiness in the bed nucleus of the stria terminalis following acute stress

Repeated acute restraint stress significantly affected the morphological properties of PNNs in the STMA, as determined by WFA labelling, confocal microscopy and image analysis (Figure 5). Stress significantly influenced the area of the individual holes within the PNN unit (Figure 5A). Specifically, the average area of the holes in the PNN matrix within the STMA was twice as large in mice following stress compared to stress-naïve animals (control; Figure 5A; stress: 0.0185 ± 0.012 vs. control: 0.0086 ± 0.0041, p < 0.0001). The perimeter of holes within the PNN matrix in the BNST was at least 35% greater in mice following stress compared to stress-naïve mice (Figure 5A; control: 0.46 ± 0.11, stress: 0.65 ± 0.20, p < 0.0001). The intensity of the ECM surrounding each hole was also altered following the restraint (Figure 5A). Specifically, the mean intensity of ECM was significantly reduced in PNNs within the BNST in mice following stress compared to control animals (Figure 5A; control: 669.9 ± 189.5, stress: 509.1 ± 206.2, p=0.0005).

**Figure 5.**
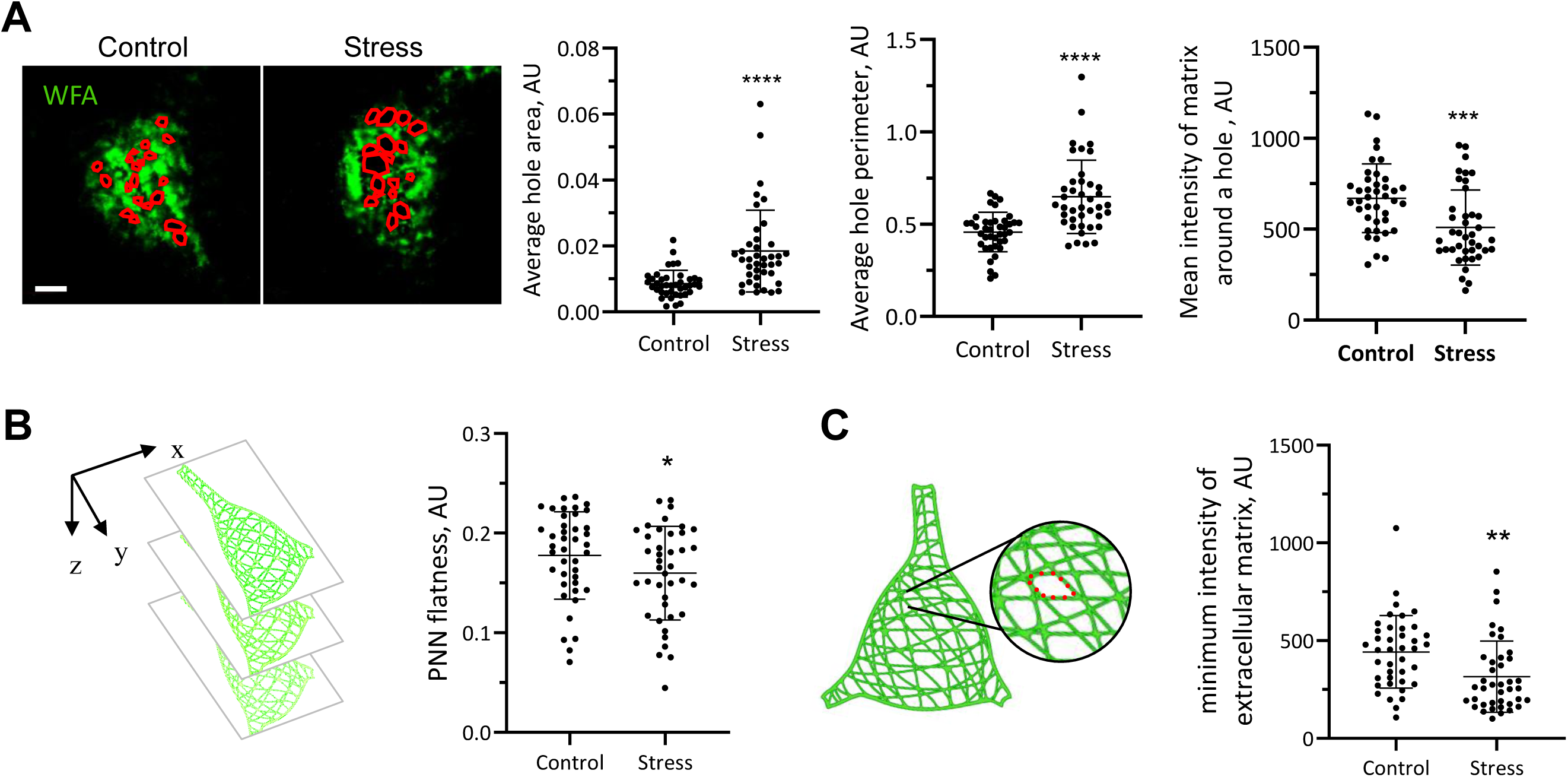
Fine morphology of perineuronal nets in the bed nucleus of the stria terminalis is vulnerable to stress. (A) Representative images of individual perineuronal net (PNN) units identified with WFA staining within the bed nucleus of the stria terminalis (BNST), medial division, anterior part (STMA) from stress naïve (control) and 7 days post-stress (stress) mice. Holes measured within the units are highlighted in red; scale bar = 5µm. The average hole area and perimeter within a BNST PNN unit is increased while the fluorescence intensity of the extracellular matrix surrounding the holes is reduced in the stress group in comparison to control mice. (B) A schematic representation of PNN flatness, determined by the standard deviation of the fluorescence intensity of the extracellular matrix across z-planes. Stress induced reduction in PNNs’ flatness. (C) A schematic representation of how PNN leakiness was determined: finding the minimum values of extracellular matrix fluorescence intensity around the perimeter of the individual holes within the PNN unit, in 3D space. The leakiness of perineuronal nets increases following stress, as evidenced by the reduction in the minimum intensity of the extracellular matrix surrounding a PNN hole. Data are presented as individual values and mean±SD; Control group: N (PNNs) = 41; stress: N (PNNs) = 40; 3 brain sections per animal, 5-6 PNNs per animal, 7 mice per group; *p<0.05, **p<0.01, ***p<0.001, ****p<0.0001, unpaired t-test. Schematic diagrams made using BioRender.

PNN flatness and leakiness were evaluated to further investigate the PNN morphology s following stress. The flatness of PNNs was determined by the standard deviation of the ECM intensity across z-planes to assess the arrangement of the PNN in 3D space (Figure 5B). PNNs in the BNST were flatter following stress compared to those in control animals (Figure 5B; stress: 0.16 ± 0.047 vs. control: 0.18 ± 0.044, p = 0.0414). The leakiness of PNNs, a measure of the structural complexity of the ECM, was determined by finding the values for the minimum intensity of the ECM around the perimeter of the PNN holes (Figure 5C). Stress increased the leakiness of PNNs in the BNST; the minimum ECM intensity of PNNs was 27% lower following stress compared to control animals (Figure 5C; stress: 315.5 ± 182 vs. control: 442.4 ± 185.8, p = 0.0028).

In summary, stress significantly affected the morphology of PNNs within the BNST. PNNs occupy less three-dimensional space and become less structurally complex, and therefore more structurally vulnerable, following stress.

### Removal of perineuronal nets in the bed nucleus of the stria terminalis exacerbates the response to repeated acute stress

Finally, we tested whether interfering with PNNs in the BNST would affect the response to acute repeated stress. First, we confirmed that a single injection of chondroitinase ABC (ChABC) resulted in a significant reduction of PNNs in the STMA, where in most animals none or only a few PNNs were detected (Figure 6A, ChABC: 3.33 ± 1.63 vs control: 27 ± 9.77, p=0.0003).

**Figure 6.**
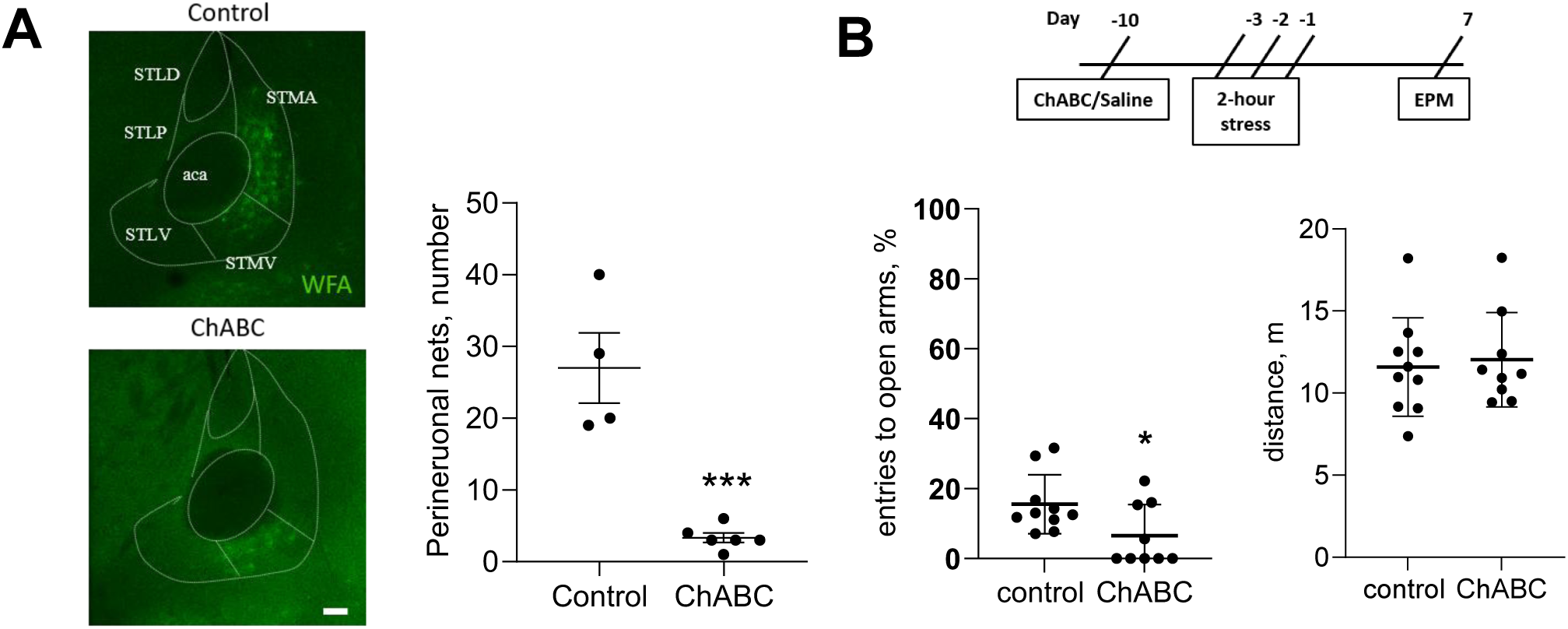
Removal of perineuronal nets in the bed nucleus of the stria terminalis exacerbates response to stress. (A) PNNs in the BNST were successfully digested and have not reinstated 14 days post ChABC injection as illustrated by significant reduction in their number in mice injected with ChABC (ChABC) compared to animals (control) that received a saline injection: representative images and quantification of WFA-labelled perineuronal nets. (B) Schematic timeline of the experiment: 7 days following injection of chondroitinase ABC (ChABC), mice were restraint for 2 hours during 3 consecutive days and then 7 days after the last restraint session tested on the elevated plus maze (EPM). PNN removal resulted in exacerbated response to acute restraint stress as measured by reduced percentage of entries into open arms on the EPM without affecting overall activity of mice as measured by unchanged distance travelled on the EPM. Data are presented as individual values and mean±SD; A: N(control)=4; N(ChABC)=6; B: N(control)= 10; N(ChABC)=9; ***p<0.001, *p<0.05, unpaired t-test.

Next, mice received a single injection of either saline or ChABC into the STMA. Following a 7 days recovery, mice were exposed to the acute restraint stress over three consecutive days, and 7 days later were tested on the EPM (Figure 6B).

PNN removal in the STMA of the BNST resulted in exacerbated response to repeated restraint stress as measured by a decreased proportion of entries into the open arms of the EPM (Figure 6B; ChABC: 6.58 ± 8.86 vs control: 15.52 ±8.4, p=0.0374) with no effect on animal activity in the maze (Figure 6B; ChABC: 12.03m ± 0.95 vs control: 11.59m ± 0.95).

## DISCUSSION

In the current study, we showed that PNNs within the BNST are first detected at P30, with the greatest expression occurring temporally at P210 and spatially in the anterior part of its medial division (STMA). BNST PNNs were sensitive to stress with PNN count decreasing following repeated acute restraint stress. The fine structure of BNST PNNs was particularly vulnerable to acute stress, exhibiting an increased area of holes within the structure and an overall increase in leakiness. Removal of BNST PNNs prior to repeated restraint stress was anxiogenic in mice indicating that BNST PNNs are required for mediating a healthy emotional response to an acute stressor.

PNNs have been previously identified in a wide variety of brain regions including sensory and motor cortices, the hippocampus, amygdala, thalamic regions and the cerebellum [24, 40, 44, 45]. However, the appearance, composition, spatiotemporal development and function of PNNs in the BNST have remained largely unknown, with only a few studies reporting their presence within the BNST until now [34–36]. In the current study of the BNST, PNNs were exclusively identified around the neuronal cell bodies and proximal dendrites of neurons (Figure 1, 2).

PNN organisation changes during development, shifting from a more granular and diffuse structure in the earlier postnatal period to a more reticular structure as animals reach adulthood [46]. Although BNST PNNs lack the regular lattice of cortical nets, their granular matrix still contains discrete gaps that can be quantified. Notably, some prefrontal regions maintain PNNs with a granular morphology in adulthood, which is hypothesised to be due to the highly plastic nature of the PFC [46]. Granular PNNs exhibit greater structural vulnerability and could therefore theoretically undergo morphological changes more readily [46, 47]. WFA labelling in the BNST revealed similarly granular PNNs (Figure 1, 2), which lacked the distinct lattice-like structure observed in PNNs in the visual cortex or hippocampus [48]. However, unlike in the PFC, WFA labelling in the BNST extended to the proximal dendrites of PNN-enwrapped neurons (Figure 1, 2). The BNST, like the PFC, is a plastic brain region, particularly in response to stress and chronic exposure to drugs, and such specific morphology of PNNs is likely to reflect BNST plasticity [49–51]. Altogether, the morphological structure of PNNs within the BNST, combined with previous evidence of neuronal plasticity in the BNST, suggests that BNST PNNs could be more prone to remodelling and/or structural damage than PNNs in other brain regions.

As key regulators of synaptic plasticity, PNNs form at the end of critical periods of high synaptic plasticity, which are particularly sensitive to experience and environmental influences [23, 24, 41, 42]. Critical periods are essential for the development of both sensory systems, such as song learning in birds or the establishment of ocular dominance, and memory systems, particularly those related to emotional memories. Critical periods in different brain regions occur at different times during postnatal development. Consequently, the time at which PNNs begin to be expressed across brain regions is also variable. In mice, PNNs are first detected in the brainstem at P4, in the somatosensory cortex and hippocampus at P14, in the amygdala at P16 and in the piriform cortex at P28 [24, 40, 44, 52]. Comparatively, WFA labelling in the BNST was first detected at P20 here, in line with the previous study by Horii-Hayashi et al (2015) [35]. However, the distinct morphological structure of mature PNNs was not detected by WFA labelling in present study until P30-P60.

Available evidence from rodent and human studies suggests that PNNs mature during adolescence, which is consistent with their developmental role in various brain regions [23, 24, 41]. In mice, PNNs in the amygdala mature around P45, whereas in the hippocampus PNNs reach maturity at P60, and in the somatosensory and piriform cortices they mature at an even later time point, P77 [24, 40, 44, 52]. In a human study investigating the expression of PNNs in postmortem brains, PNNs were shown to mature in the hippocampus and medial PFC around 14 years of age [53].

We also investigated PNN number and morphology within the BNST at P210 and P365 which roughly corresponds to the ‘mature adult’ and ‘middle age’ stages, respectively [25] as the BNST is known to continue developing into adulthood [37, 54]. The comparatively late development of PNNs in the BNST, up to P60 in some subnuclei and beyond this time point in others, suggests that any potential critical period facilitated by BNST is likely to be longer than those studied in other brain regions [23, 24, 41]. Conversely, PNNs in the BNST may not be responsible for the regulation of a critical period. As PNNs formation are experience-driven, their structure and accumulation in the BNST may develop as a result of gained experiences, most likely those of an emotional nature.

PNNs were quantified using two distinct methods, WFA labelling intensity and PNN count, both confirming an increase in PNN expression throughout development in the BNST. WFA labelling of the BNST at different developmental time points revealed specific subnuclear regions of high-density PNN expression; particularly the STMA (Figure 2; Supplementary Tables 1-8). The STMA is one of the lesser studied BNST subnuclei, but the presently established connections between the STMA and other brain regions suggest that STMA could mediate the anxiogenic effects of the BNST [17]. The STMA receives olfactory projections from the medial amygdala, contextual projections from the subiculum and dense glutamatergic projections from the basomedial amygdala [16, 18]. Given that threatening odours and contexts can generate an anxiogenic response, evidence suggests the STMA could, in part, may mediate the anxiogenic response to such stimuli.

We used a repeated restraint stress model, three consecutive days of two-hour sessions, to induce anxiety-like behaviour in mice. This was enough to produce anxiety-like behaviour in the EPM, shown by a reduced proportion of open arm entries (Figure 4B). Stressed mice also showed reduced locomotor activity (Figure 4C) which has been also previously reported in stress paradigms. Decreased exploration after stress has been viewed both as a methodological confound, reflecting sedation, malaise or locomotor incapacity [55–57], and as sign of a negative emotional state [57, 58]. Importantly, our main measure of anxiety-like behaviour, the proportion of entries made into the open arms, is expressed relative to total arm entries. It therefore partly controls for overall differences in activity, so the reduction in open-arm exploration is unlikely to be explained by hypolocomotion alone.

We hypothesised that PNNs in the anteromedial BNST provide structural stability to emotional circuits, so that intact nets limit stress-induced anxiety while their degradation weakens this protection. Several observations support our hypothesis. To begin with, PNN density was sensitive to stress: increased anxiety-like behaviour was accompanied by fewer PNNs in the BNST (Figure 4E). In addition, the integrity of the remaining nets was compromised. PNNs in the STMA showed greater hole area and perimeter, reduced intensity of the matrix surrounding the holes, and greater leakiness, measured as a reduced minimum intensity of the ECM around the holes (Figure 5). In contrast, the variability of PNN fluorescence across z-planes was unchanged. These results indicate that, following stress, PNNs in the STMA occupy three-dimensional space similar to control animals, however their organisation becomes much less structurally complex and therefore more structurally vulnerable. Lastly, PNNs were necessary for limiting the behavioural impact of stress: removing STMA PNNs with ChABC before restraint further reduced open arm entries in the EPM. Together, these findings indicate that both the number and the structural integrity of BNST PNNs are reduced by stress, and that their presence is necessary for limiting stress-induced anxiety-like behaviour.

Our findings contrast with some reports of stress-induced increases in PNNs in other limbic areas. For example, 24-hour restraint stress increases PNN density in the basolateral amygdala [31], while chronic stress increases the amount of prefrontal PNNs [32]. Conversely, early-life adversity reduces PNN intensity in the basolateral amygdala, and removing these PNNs enhances innate fear responses [59]. The divergent effects of stress on PNNs across studies may reflect differences in brain region, the duration of stress, the time between stress and tissue collection, and the PNN markers used. If PNN loss is followed by compensatory rebuilding, studies may also be capturing different phases of the same remodelling process. Rather than uniformly degrading PNNs, stress may therefore change them in either direction depending on region and timing, with loss and loosening of the nets being the main effect in the BNST under our conditions.

Stress-induced ECM remodelling may involve various mechanisms including proteolysis by matrix metalloproteinases and ADAMTS proteases or microglia-dependent engulfment of the ECM [60–62]. Degradation would first loosen the fine structure of existing nets, increasing hole size and leakiness, and with continued stress leading to the loss of whole nets. Reduced PNN integrity could then reopen plasticity in STMA neurons. This could happen through destabilised synaptic inputs, altered excitability, or reduced perisomatic inhibition, making STMA output more readily shaped by stress-related inputs from the amygdala and ventral subiculum. Given the STMA’s connections with these regions, such a change could bias BNST output towards heightened anxiety.

Our findings are relevant to the actions of pharmacological treatments shaping emotional recovery and mood, with effects depending on drug class, region and stress state. Chronic treatment with a specific serotonin reuptake inhibitor fluoxetine reduces PNN number in the PFC and hippocampus [63], and serotonin-norepinephrine reuptake inhibitor venlafaxine promotes PNN degradation through MMP-9 [60], effects thought to reopen plasticity that can support fear extinction and recovery from stress. Consistent with this, fluoxetine combined with extinction training produces long-lasting fear erasure in adult mice, an effect not seen with either treatment alone, and SSRI treatment reopens juvenile-like plasticity in the adult visual cortex [64, 65]. Chronic fluoxetine also alters cortical interneuron structure, connectivity and plasticity in a manner resembling critical-period plasticity [66]. Rapid acting antidepressant ketamine, by contrast, triggers microglial disassembly of cortical PNNs [62], whereas in chronically stressed mice ketamine preserves PNNs by suppressing microglial activation [67]. PNN removal in the ventral hippocampus abolishes ketamine’s sustained antidepressant effect [68]. PNN integrity therefore appears necessary for some lasting therapeutic effects rather than being uniformly degraded by treatment. Our results suggest that reduced PNN integrity is not beneficial in itself. Rather, it may open a window in which circuit function becomes more strongly shaped by ongoing experience, worsening the effects of stress when PNNs are degraded during stress exposure, but potentially enabling recovery when paired with treatment or a safe environment. Whether such treatments alter PNNs in the BNST, and whether this contributes to their effects on anxiety, remains to be defined.

Several limitations of the present study should be considered. Stress produced two concurrent changes in BNST PNNs: fewer nets and remodelling of the remaining nets’ fine structure, with larger hole area and perimeter and greater leakiness. Our design cannot separate these two changes, so it remains unclear whether the anxiogenic effect of stress depends mainly on PNN loss, on remodelling of nets that persist, or on both. The distinction is important, since a complete loss of PNNs and a partial loosening of their structure would be expected to affect synaptic stability and plasticity differently. Moreover, ChABC degrades chondroitin sulphate proteoglycans non-selectively, leading to a near-complete loss of PNNs in the targeted region. Our ChABC experiment therefore tests whether PNN presence is necessary for limiting stress-induced anxiety-like behaviour. It does not directly test the contribution of the subtler, stress-induced reorganisation of PNN architecture. Future studies could separate these processes in two ways, for example by blocking the endogenous mechanisms of PNN remodelling during stress, such as matrix metalloproteinase activity or microglial engulfment. A further limitation is the use of WFA as the only marker of PNNs. WFA recognises specific chondroitin sulphate glycosylation patterns and does not detect all PNNs, so WFA-negative nets, such as those containing aggrecan but lacking the WFA-binding epitope, will have been missed. therefore,s may therefore underestimate the total PNN population, and stress-induced reductions in WFA labelling may partly reflect changes in glycosylation rather than loss of the whole net. Finally, all experiments were carried out in male mice only. Anxiety disorders are more prevalent in women than in men, yet preclinical studies of anxiety have historically excluded female rodents [69]. The BNST is one of the most sexually dimorphic regions of the rodent brain, and its development continues into adulthood in a sex-dependent manner [37]. Notably, stress produces sex-specific and long-lasting changes in BNST neuronal activity, with some effects persisting in females but not males [69]. The spatiotemporal pattern of BNST PNN development, their sensitivity to stress, and their role in anxiety-like behaviour may therefore differ in females, so our findings should not be generalised across sexes without direct testing.

Taken together, we uncovered that PNNs within the BNST are sensitive to stress: repeated restraint reduces their number and degrades their fine structure. Removing BNST PNNs before repeated restraint stress further increased anxiety-like behaviour. This indicates that BNST PNNs are necessary for limiting the anxiogenic consequences of restraint stress and may contribute to a healthy emotional response to stressful experiences.

## Data availability statement

Data will be made available upon request to authors

## Funding statement

The financial support was provided by AMS Springboard award SBF005\1102 and the MRC Career Development Award MR/T031115/1 to VM, GW4 BioMed2 MRC Doctoral Training Partnership, funded by MRC/UKRI MR/W006308/1 to KM/VM, the Marie Sklodowska Curie ITN “Extrabrain” ID 606950, The LeverhulmeTrust and the Cleopatra Trust grants to RP.

## Conflicts of interest

The authors report no conflicts of interest.

## Notes

### Competing Interest Statement

The authors have declared no competing interest.

