## supplementary tables for "Perineuronal nets in the bed nucleus of stria terminalis mediate stress-induced anxiety behaviour in adult male mice"

| Tukey's multiple comparisons test | Significant? | Summary | Adjusted P Value |
|---|---|---|---|
| STLV vs. STMV | No | ns | 0.9733 |
| STLV vs. STMA | No | ns | 0.9996 |
| STLV vs. STLP | No | ns | 0.8930 |
| STLV vs. STLD | No | ns | 0.1181 |
| STMV vs. STMA | No | ns | 0.9272 |
| STMV vs. STLP | No | ns | 0.9981 |
| STMV vs. STLD | No | ns | 0.3234 |
| STMA vs. STLP | No | ns | 0.8037 |
| STMA vs. STLD | No | ns | 0.0806 |
| STLP vs. STLD | No | ns | 0.4834 |

**Supplementary Table 1. Statistics for comparisons of perineuronal net number in the anterior subnuclei of the bed nucleus of the stria terminalis at P30.**

| Tukey's multiple comparisons test | Significant? | Summary | Adjusted P Value |
|---|---|---|---|
| STLV vs. STMV | No | ns | 0.9997 |
| STLV vs. STMA | No | ns | 0.0906 |
| STLV vs. STLP | No | ns | 0.9909 |
| STLV vs. STLD | No | ns | 0.9988 |
| STMV vs. STMA | No | ns | 0.1288 |
| STMV vs. STLP | No | ns | 0.9673 |
| STMV vs. STLD | No | ns | 0.9907 |
| STMA vs. STLP | Yes | * | 0.0366 |
| STMA vs. STLD | No | ns | 0.0535 |
| STLP vs. STLD | No | ns | 0.9997 |

**Supplementary Table 2. Statistics for comparisons of perineuronal net number in the anterior subnuclei of the bed nucleus of the stria terminalis at P60.**

| Tukey's multiple comparisons test | Significant? | Summary | Adjusted P Value |
|---|---|---|---|
| STLV vs. STMV | No | ns | 0.9657 |
| STLV vs. STMA | Yes | **** | <0.0001 |
| STLV vs. STLP | No | ns | 0.9838 |
| STLV vs. STLD | No | ns | 0.9917 |
| STMV vs. STMA | Yes | **** | <0.0001 |
| STMV vs. STLP | No | ns | >0.9999 |
| STMV vs. STLD | No | ns | 0.9996 |
| STMA vs. STLP | Yes | **** | <0.0001 |
| STMA vs. STLD | Yes | **** | <0.0001 |
| STLP vs. STLD | No | ns | >0.9999 |

**Supplementary Table 3. Statistics for comparisons of perineuronal net number in the anterior subnuclei of the bed nucleus of the stria terminalis at P210.**

| Tukey's multiple comparisons test | Significant? | Summary | Adjusted P Value |
|---|---|---|---|
| STLV vs. STMV | Yes | * | 0.0426 |
| STLV vs. STMA | Yes | **** | <0.0001 |
| STLV vs. STLP | No | ns | 0.7316 |
| STLV vs. STLD | No | ns | 0.9646 |
| STMV vs. STMA | Yes | **** | <0.0001 |
| STMV vs. STLP | No | ns | 0.4092 |
| STMV vs. STLD | Yes | ** | 0.0094 |
| STMA vs. STLP | Yes | **** | <0.0001 |
| STMA vs. STLD | Yes | **** | <0.0001 |
| STLP vs. STLD | No | ns | 0.3496 |

**Supplementary Table 4. Statistics for comparisons of perineuronal net number in the anterior subnuclei of the bed nucleus of the stria terminalis at P365.**

| Tukey's multiple comparisons test | Significant? | Summary | Adjusted P Value |
|---|---|---|---|
| STLV vs. STMV | No | ns | >0.9999 |
| STLV vs. STMA | No | ns | 0.4432 |
| STLV vs. STLP | No | ns | 0.9788 |
| STLV vs. STLD | No | ns | 0.9179 |
| STMV vs. STMA | No | ns | 0.5101 |
| STMV vs. STLP | No | ns | 0.9585 |
| STMV vs. STLD | No | ns | 0.8752 |
| STMA vs. STLP | No | ns | 0.1901 |
| STMA vs. STLD | No | ns | 0.1181 |
| STLP vs. STLD | No | ns | 0.9988 |

**Supplementary Table 5. Statistics for comparisons of perineuronal net intensity in the anterior subnuclei of the bed nucleus of the stria terminalis at P30.**

| Tukey's multiple comparisons test | Significant? | Summary | Adjusted P Value |
|---|---|---|---|
| STLV vs. STMV | No | ns | 0.9106 |
| STLV vs. STMA | Yes | * | 0.0179 |
| STLV vs. STLP | No | ns | 0.5793 |
| STLV vs. STLD | No | ns | 0.9831 |
| STMV vs. STMA | Yes | ** | 0.0027 |
| STMV vs. STLP | No | ns | 0.9668 |
| STMV vs. STLD | No | ns | 0.9975 |
| STMA vs. STLP | Yes | *** | 0.0006 |
| STMA vs. STLD | Yes | ** | 0.0055 |
| STLP vs. STLD | No | ns | 0.8690 |

**Appendix Table 6. Statistics for comparisons of perineuronal net intensity in the anterior subnuclei of the bed nucleus of the stria terminalis at P60.**

| Tukey's multiple comparisons test | Significant? | Summary | Adjusted P Value |
|---|---|---|---|
| STLV vs. STMV | No | ns | 0.3283 |
| STLV vs. STMA | Yes | **** | <0.0001 |
| STLV vs. STLP | Yes | ** | 0.0011 |
| STLV vs. STLD | Yes | * | 0.0102 |
| STMV vs. STMA | Yes | **** | <0.0001 |
| STMV vs. STLP | No | ns | 0.0981 |
| STMV vs. STLD | No | ns | 0.4515 |
| STMA vs. STLP | Yes | **** | <0.0001 |
| STMA vs. STLD | Yes | **** | <0.0001 |
| STLP vs. STLD | No | ns | 0.8891 |

**Appendix Table 7. Statistics for comparisons of perineuronal net intensity in the anterior subnuclei of the bed nucleus of the stria terminalis at P210.**

| Tukey's multiple comparisons test | Significant? | Summary | Adjusted P Value |
|---|---|---|---|
| STLV vs. STMV | No | ns | 0.2773 |
| STLV vs. STMA | Yes | *** | 0.0004 |
| STLV vs. STLP | No | ns | 0.4885 |
| STLV vs. STLD | No | ns | 0.8429 |
| STMV vs. STMA | Yes | **** | <0.0001 |
| STMV vs. STLP | No | ns | 0.9942 |
| STMV vs. STLD | No | ns | 0.8450 |
| STMA vs. STLP | Yes | **** | <0.0001 |
| STMA vs. STLD | Yes | **** | <0.0001 |
| STLP vs. STLD | No | ns | 0.9712 |

**Appendix Table 8. Statistics for comparisons of perineuronal net intensity in the anterior subnuclei of the bed nucleus of the stria terminalis at P365.**
